# Rapid and Inexpensive Whole-Genome Sequencing of SARS-CoV-2 using 1200 bp Tiled Amplicons and Oxford Nanopore Rapid Barcoding

**DOI:** 10.1101/2020.05.28.122648

**Authors:** Nikki E. Freed, Markéta Vlková, Muhammad B. Faisal, Olin K. Silander

**Affiliations:** School of Natural and Computational Sciences, Massey University, Auckland, New Zealand

## Abstract

Rapid and cost-efficient whole-genome sequencing of SARS-CoV-2, the virus that causes COVID-19, is critical for understanding viral transmission dynamics. Here we show that using a new multiplexed set of primers in conjunction with the Oxford Nanopore Rapid Barcode library kit allows for faster, simpler, and less expensive SARS-CoV-2 genome sequencing. This primer set results in amplicons that exhibit lower levels of variation in coverage compared to other commonly used primer sets. Using five SARS-CoV-2 patient samples with C_q_ values between 20 and 31, we show that high-quality genomes can be generated with as few as 10,000 reads (approximately 5 Mbp of sequence data). We also show that mis-classification of barcodes, which may be more likely when using the Oxford Nanopore Rapid Barcode library prep, is unlikely to cause problems in variant calling. This method reduces the time from RNA to genome sequence by more than half compared to the more standard ligation-based Oxford Nanopore library preparation method at considerably lower costs.

## Introduction

The earliest known outbreak of COVID-19 occurred in Wuhan, China in late 2019. By May of 2020, the disease had spread to more than 200 countries and territories, with more than five million total confirmed cases (Johns Hopkins Coronavirus Resource Center). To slow the spread of this disease, it is crucial to understand the origin and dynamics of outbreak events. Genome sequencing of SARS-CoV-2, the causative agent of this disease, offers an effective means to infer transmission events, and such genomic surveillance has been used successfully to ascertain transmission events even in the absence of additional epidemiological data (Seemann et al. 2020).

Several approaches have been used to sequence the SARS-CoV-2 genome, including metagenomics (Manning et al. 2020), sequence capture (Gohl et al. 2020), SISPA (Moore et al. 2020), and multiplex PCR (Gohl et al. 2020; Itokawa et al. 2020; Resende et al. 2020; Moore et al. 2020; Eden et al. 2020), followed by next generation sequencing using either the Illumina or Oxford Nanopore platforms. Due to its simplicity and economy, using multiplexed PCR amplicons is perhaps the most common approach. This technique has been used to successfully sequence thousands of genomes over the first few months of the COVID-19 outbreak (GISAID Initiative). However, multiplexed amplicon sets often exhibit uneven amplification across the genome, with up to 100-fold differences in the concentration of different amplicons (Itokawa et al. 2020). In addition, the most common methods of library preparation for next-generation sequencing remain relatively expensive, even when samples are multiplexed.

To alleviate these problems, here we describe an approach using multiplexed 1200 base pair (bp) tiled amplicons with the Oxford Nanopore Rapid Barcoding kit (SQK-RBK004). Briefly, two PCR reactions are performed for each SARS-CoV-2 positive patient sample to be sequenced. One PCR reaction contains thirty primers that generate the odd numbered amplicons (‘Pool 1’), while the second PCR reaction contains twenty eight primers that generate the even numbered amplicons (‘Pool 2’; **Fig. 1**). After PCR, the two amplicon pools are combined and can be used for a range of downstream sequencing approaches. Here we use the Oxford Nanopore Rapid barcoding kit which enables relatively easy library preparation of these amplicons to achieve rapid, simple, and cost-effective sequencing.

**Figure 1.**
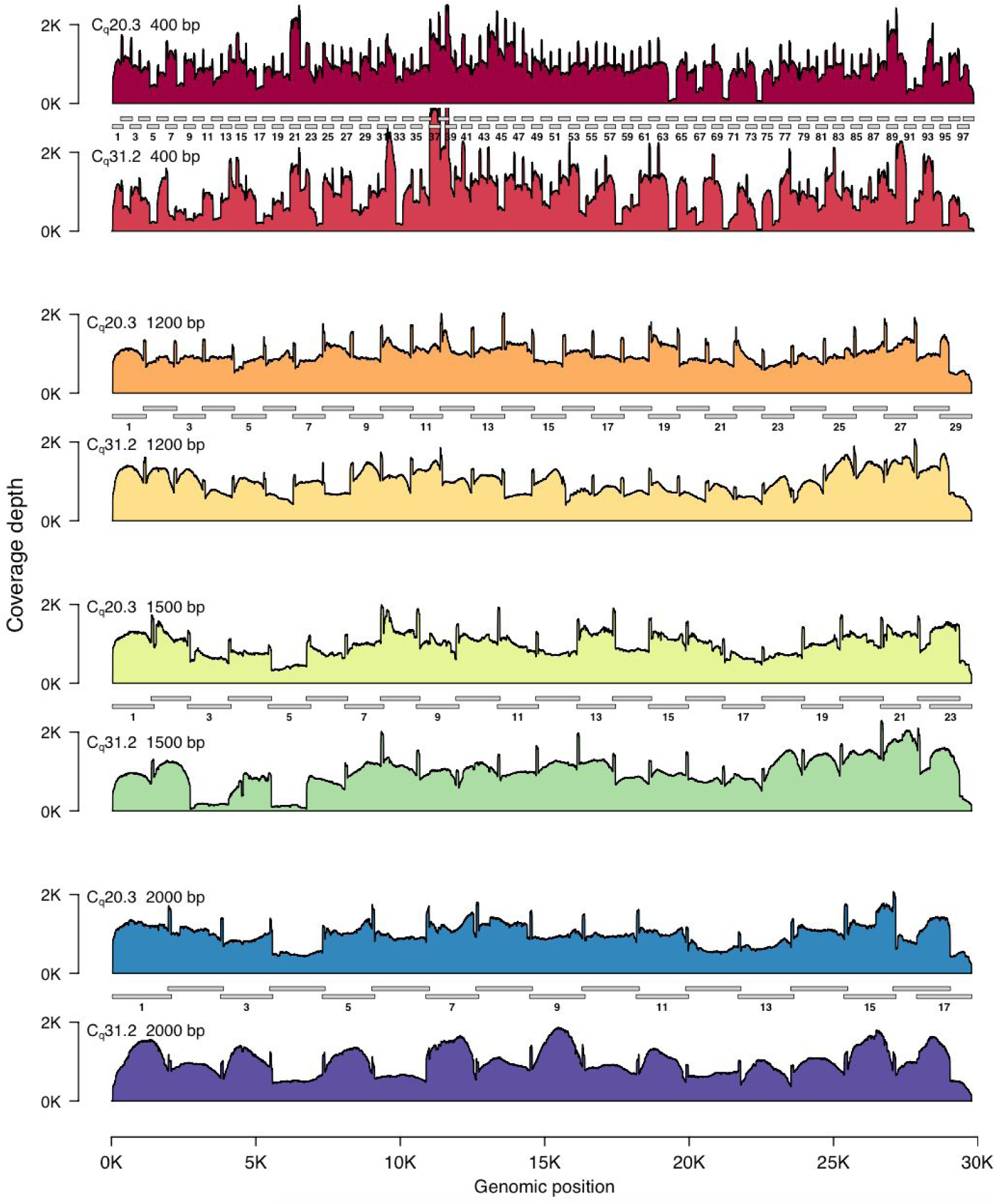
SARS-CoV2 genome coverage plots for different amplicon sets. We performed amplicon sequencing of the SARS-CoV-2 genome with amplicons ranging from 400 bp (top) to 2000 bp in size (bottom). Amplicon sets are shown as grey bars, with the amplicons in ‘Pool 1’ numbered (see Methods). Read coverage is scaled so that mean coverage is 1000X for all amplicon sets. For each set of amplicons we sequenced a low C_q_ sample (20.3) and a high C_q_ sample (31.2). Each amplicon set is shown in pairs. The upper plot is the C_q_ 20.3 sample; the lower plot is the C_q_ 31.2 sample. The 1200 bp and 2000 bp amplicon sets exhibit relatively even coverage across the entire SARS-CoV-2 genome. However, note that for the high C_q_ 2000 bp amplicon set, all amplicons in ‘Pool 1’ are approximately 1.5-fold higher levels than those in ‘Pool 2’. In contrast to the 1200 bp and 2000 bp amplicon sets, several dropout regions are apparent in the 400 bp and 1500 bp amplicon sets. In all cases, the variation in genome coverage is higher for the sample with higher C_q_.

## Materials and Methods

### RNA isolation and RT-qPCR

Five de-identified samples that had been assessed as positive by other methods were obtained from the New Zealand Auckland District Health Board. 80 µl of Viral Transport Media that had previously stored a nasopharyngeal swab from a patient infected with SARS-CoV-2 were used for RNA isolation using the QIAamp Viral RNA Mini spin kit (Qiagen, Cat No./ID: 52904) according to manufacturer specifications, with the following modifications: sample volume was brought up from 80 µl to 140 µl using 1x PBS, and RNA was eluted using two elutions with 40 µl Buffer AVE for a final volume of approximately 80 µl.

To determine C_q_ values, we performed quantitative RT PCR. We used the 2x SensiFAST Probe No-ROX One-Step Mix (Bioline, Cat No./ID: BIO-76005), according to manufacturer specifications using 2.5 µl RNA template. Primers and probes were from the CDC 2019-nCoV CDC Assay (IDT, Cat No./ID: 10006606) targeting the N1 and N2 regions of the nucleocapsid phosphoprotein of SARS-CoV-2. NP targets the RNase P gene for detection of human nucleic acids and serves as an internal control for sample integrity. Reactions were run at a total volume of 20 µl on the Liberty16 mobile real-time PCR device (Ubiquitome). The results are shown in **Table 2**.

**Table 1.** Sample C_q_ values. qPCR was performed using the CDC Primer probe set (IDT, Cat No./ID: 10006606). N1 and N2 target the nucleocapsid phosphoprotein (N gene) of SARS-CoV2. NP targets the RNase P gene for detection of human nucleic acids and serves as a control for sample integrity.

| Sample | N1 | N2 | NP |
| --- | --- | --- | --- |
| AKL-MU-1 | 20.3 | 20.0 | 27.4 |
| AKL-MU-2 | 20.6 | 20.4 | 27.0 |
| AKL-MU-3 | 25.7 | 25.4 | 28.2 |
| AKL-MU-4 | 28.5 | 28.2 | 29.6 |
| AKL-MU-5 | 31.2 | 30.5 | 33.4 |

**Table 2.**
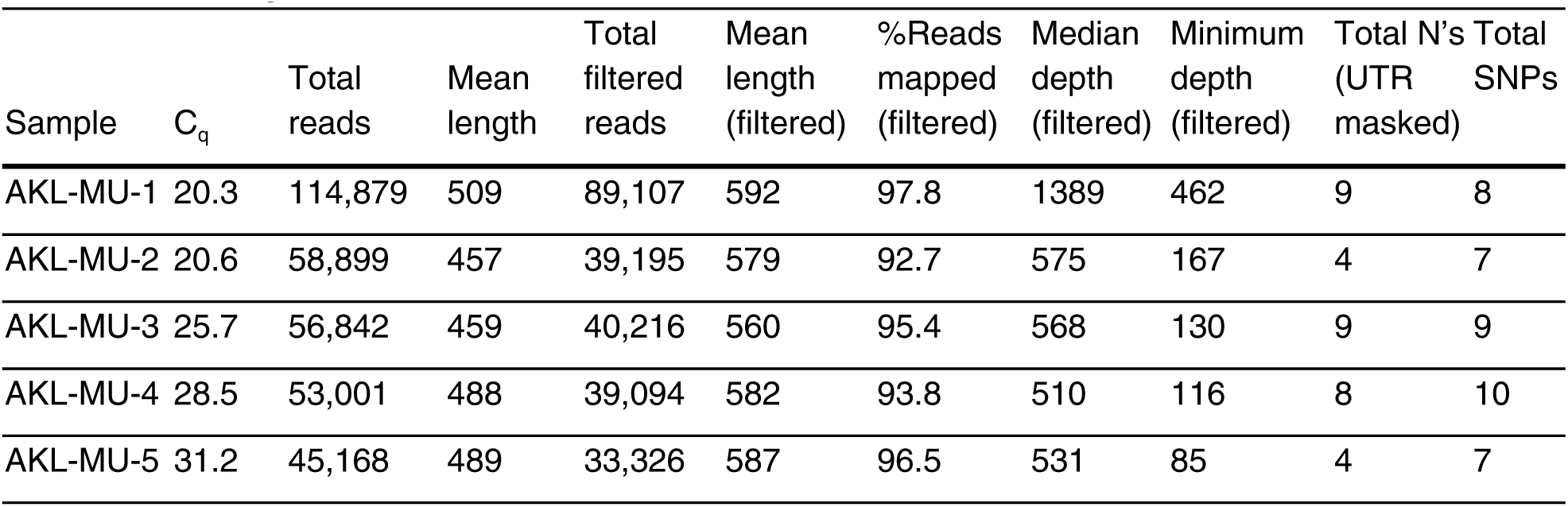
Sequencing, mapping, and assembly statistics for a four hour MinION run. The filtering steps remove reads shorter than 250 bp and longer than 1500 bp. C_q_ indicates the value for the N1 gene. The values for the N2 gene are similar.

### Whole Genome Sequencing

The detailed methods for Reverse Transcription, PCR, and library preparation have been posted to protocols.io:

https://www.protocols.io/view/ncov-2019-sequencing-protocol-rapid-barcoding-1200-bgggjttw

Below we briefly summarize the method and provide additional notes on the genome assembly.

Two separate PCR reactions are done for each SARS-CoV2 positive sample. Two pools of primers are made: ‘Pool 1’ contains thirty primers that generate the odd numbered tiled amplicons, while ‘Pool 2’ contains twenty eight primers that generate the even numbered tiled amplicons for the 1200 bp set (**Fig. 1**). This tiled approach is necessary to minimise overlap between amplicons, which would decrease PCR efficiency as overlapping amplicons would anneal to each other. After the PCR is completed, we combine the two pools.

### Basecalling, assembly, and variant calling

We used Guppy Version 3.6.0 for basecalling and demultiplexing all runs. We used the ARTIC Network bioinformatics protocol for all genome assembly and variant calling steps with the 1200 bp amplicon sets (https://artic.network/ncov-2019/ncov2019-bioinformatics-sop.html), with the .bed and .tsv files adjusted accordingly to accommodate the different primer sequences and binding locations. We used 250 bp as the minimum read length cut-off and 1500 bp as the maximum length cut-off, which resulted in between 20% and 30% of the reads being filtered. The primer sequences and .bed and .tsv files necessary for the ARTIC variant calling pipeline are available on Zenodo https://zenodo.org/record/3897530#.Xuk7oGpLjep.

To quantify the number of ambiguous bases in the genome assemblies, we excluded the first and last 180 bp of the genome, which are outside of any amplicon set or which contain primer binding sites, and are thus usually masked during the ARTIC bioinformatics pipeline.

### Read subsampling

To calculate how sequencing depth affected genome coverage for each amplicon set, we first calculated the total number of Mbp collected for each amplicon set. We then scaled that total to the required sequencing depth. For example, if a total of 40 Mbp of data was collected for one amplicon set, we calculated genome coverage for that amplicon set, and then multiplied the coverage values by the relevant factor (e.g. to calculate coverage for 20 Mbp of data, we multiplied by 0.5). This removes the complicating issue that different amplicon sets have different distributions of read lengths.

For read subsampling we used seqtk (https://github.com/lh3/seqtk). To simulate read contamination, we used seqtk to subsample using a random seed, and concatenated the subsampled read sets from two different samples, performing this for all possible pairwise combinations. We then used the ARTIC Network bioinformatics pipeline after this subsampling to call variants. The steps in the pipeline include filtering to exclude short reads and likely chimeric reads, which we performed as noted above.

## Results

Our first goal was to select an amplicon set resulting in even coverage across the SARS-CoV-2 genome, to ensure that we could obtain high-quality, complete genomes with minimal sequencing depth. We tested four different tiled, multiplexed amplicon sets: the current ARTIC network 400 base pair V3 primer pool, available from Integrated DNA technology (IDT catalog number: 10006788), and three sets designed using Primal Scheme (Quick et al. 2017). These sets generate amplicons that are approximately 1200 bp, 1500 bp, and 2000 bp in size (**File S1**). In all cases, primers were multiplexed into two pools (‘Pool 1’ and ‘Pool 2’), creating a tiled amplification of the entire SARS-CoV2 genome (**Fig. 1**).

Our second goal was to reduce the time and cost of library preparation. To this end, we used the Oxford Nanopore Rapid Barcoding kit. This kit uses a transposase to attach barcodes and motor proteins to DNA molecules, with library preparation taking less than 25 minutes of hands on time and requiring minimal reagents (**Table S1**). This contrasts with the Oxford Nanopore Ligation Sequencing kit (LSK-109) that is most commonly used (J. Quick, protocols.io), which requires over two hours of hands on time for library preparation, as well as several third-party reagents. In addition to using the Rapid Barcoding kit approach, we omit a bead-based cleanup step after PCR, further reducing the time and cost.

We tested all four primer sets on two RT-qPCR positive patient samples, one with a cycle quantification value (C_q_) of 20.3, and the second with a C_q_ of 31.2 (**Table 1**), which is similar to the mean C_q_ found in sputum, feces, and pharyngeal swabs, and considerably higher than that found in nasal swabs (Wang et al. 2020). We found that all four primers sets produced adequate results. However, the 1200 bp set exhibited the least variability, especially in the high C_q_ sample (**Fig. 1**).

We quantified this variation by testing how many Mbp of read data was required to achieve 30X coverage at every position in the genome (here we use Mbp of sequencing data and not read number to mitigate the effects of differences in average read length between the samples). We found that for the 1200 bp and 2000 bp amplicon sets, 30X read coverage across the 99.9% of the genome could be achieved with only 3Mbp of data (**Fig. 2**). In contrast, for the 400 bp amplicon set, genome coverage was far more variable, and for the high C_q_ sample, even with 20Mbp of data, 30X coverage was achieved for only 99% of the genome. We thus selected the 1200bp primer set for additional experiments, as this amplicon size yielded the most consistent results.

**Figure 2.**
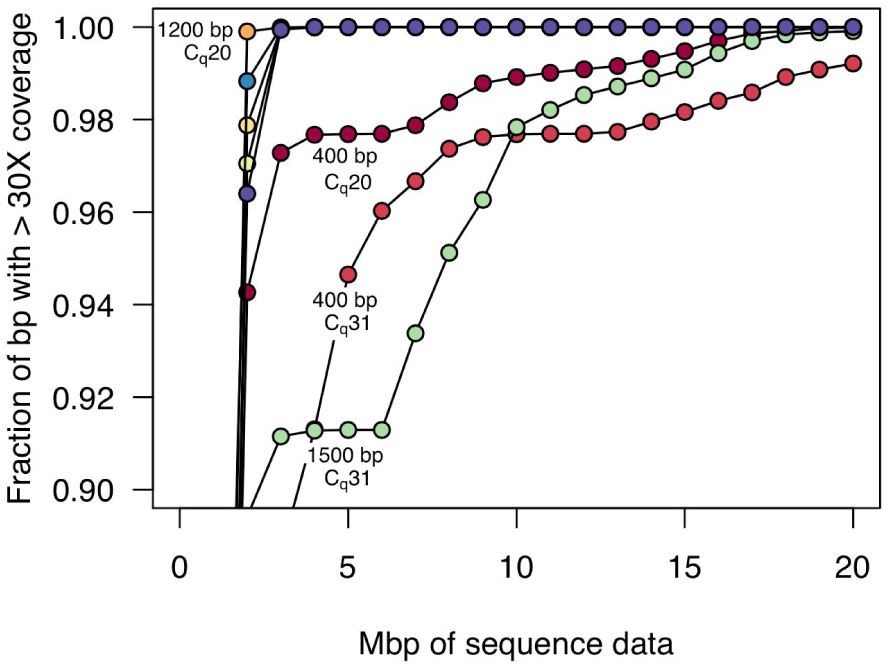
Amount of sequence data required for 30-fold genome coverage. As more sequencing data is collected, a greater fraction of the genome is covered. Here, we plot the amount of data required for 30X coverage, which is similar to the minimum level required for accurate variant calling. For both the high and low C_q_ samples, the 1200 bp and 2000 bp amplicon sets achieved more than 99.9% genome coverage with only 3 Mbp of data, and in the low C_q_ sample, the 1200 bp amplicon set achieved 99.9% coverage with only 2 Mbp of data. In contrast, the 400 bp and 1500 bp amplicon sets were more variable in coverage, especially for the high C_q_ sample. In the case of the 400 bp amplicon set, 99% genome coverage at 30X required 19 Mbp of sequence data, and 99.9% was only achieved with 33 Mbp of sequence data.

We next tested the reliability of the 1200 bp amplicon set with the Rapid Barcode kit library prep using three additional SARS-CoV-2 positive patient samples. Across these five samples, C_q_ values ranged from 20 to 31 (**Table 1**). We multiplexed all five samples on a single flow cell for four hours. Again we found relatively even genome coverage for all five samples, although for higher C_q_ samples, there again appeared to be greater variability across the genome (**Fig. 3**).

**Figure 3.**
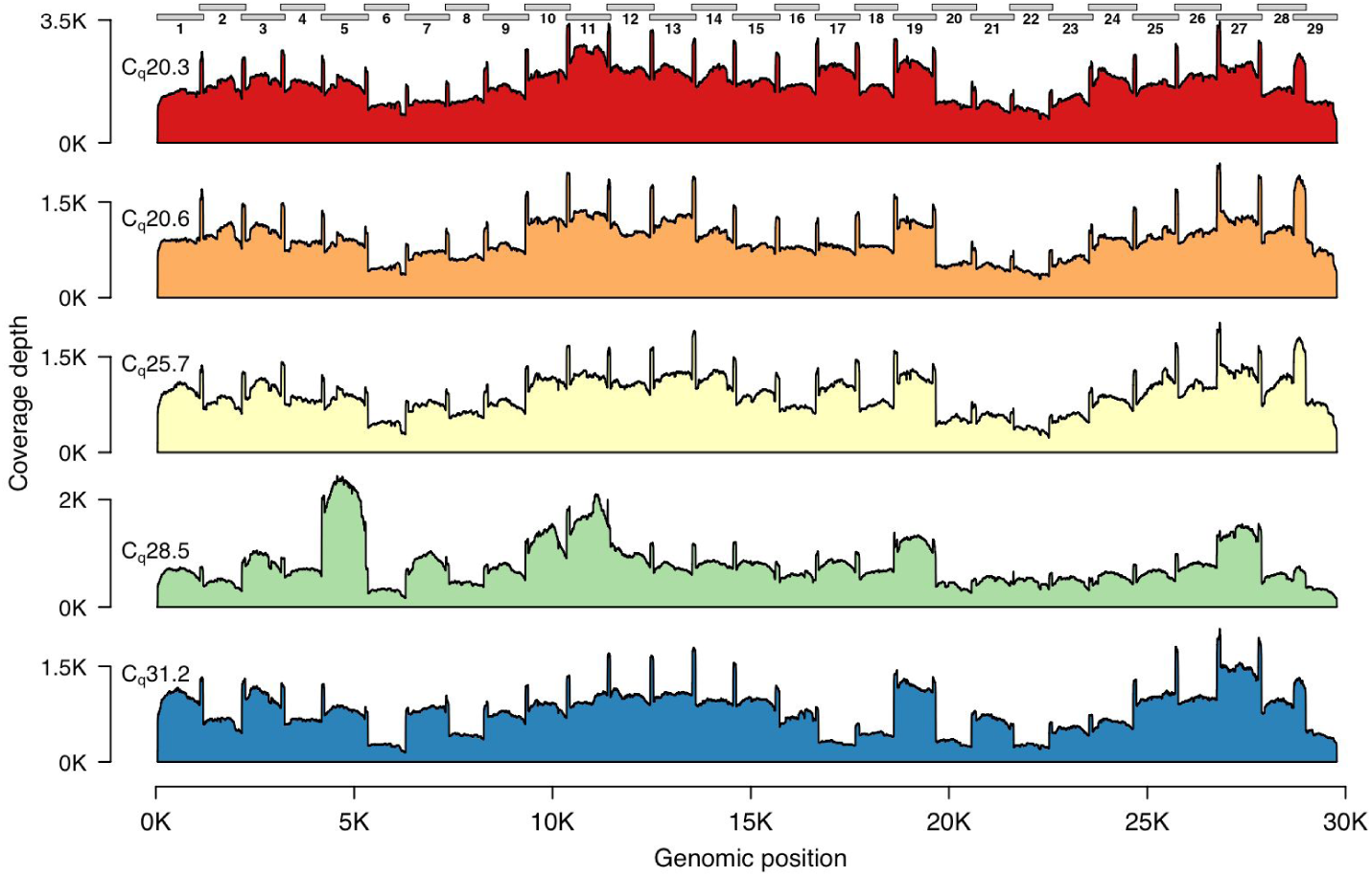
Genome coverage plots for patient samples varying in C_q_ values. The plots indicate the genome coverage for the 1200 bp amplicon set for samples with C_q_ values ranging from 20.3 to 31.2. For all samples, minimum coverage exceeds 50 at all genomic positions (excluding the 5’ and 3’ UTR). Note that the scale of the y-axes varies between plots. The locations of the amplicons are indicated above the first plot.

We used the ARTIC Network bioinformatics pipeline (see **Methods**) for genome assembly and variant calling for these samples, and in all cases obtained assemblies with fewer than 10 ambiguous base pairs (**Table 2**), excluding the 5’ and 3’ ends of the genome, which are either not part of an amplicon or which are primer sequences.

We again quantified the sequencing depth (number of reads) required to achieve sufficient genome coverage for assembly and variant calling. We downsampled the read sets for each sample, varying the number of reads from 2,500 to 30,000 (approximately 1.25 Mbp to 15 Mbp, assuming a pre-filtered average read length of 500 bp (**Table 2**)), and quantified the fraction of the genome covered at varying depths (**Fig. 4**). We found that with low C_q_ samples, very few reads were required to achieve high coverage depth across the genome. For the lowest C_q_ samples, only 10,000 reads (approximately 5 Mbp) were required to achieve 50X coverage depth over more than 99.9% of the genome.

**Figure 4.**
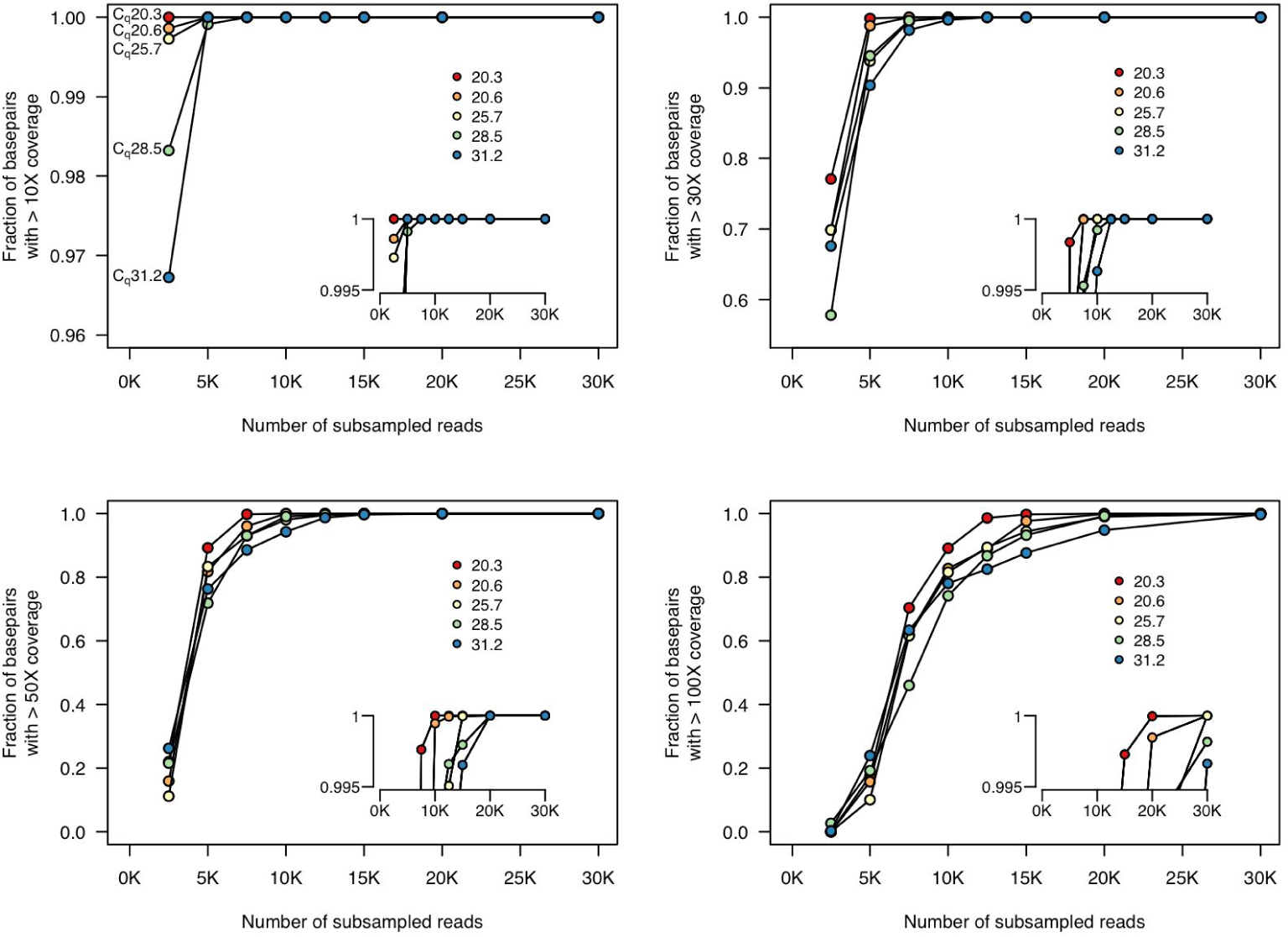
Fraction of genome covered at different sequencing depths. We subsampled from the complete set of unfiltered reads and mapped these reads to the reference sequence. For all five samples, 30X coverage of all genomic positions is achieved with only 12.5K reads. 50X coverage at all genomic positions is achieved with fewer than 20K reads. Insets show genome coverage levels at the top end of the y-axis (range from 0.995 to 1). Each line indicates the coverage for one sample. Insets show higher resolution at the upper limit of the y-axis. The colours of each sample on these plots are the same as those in **Fig. 3**. Note that the scale of the y-axis in the top left plot differs from the others.

Finally, we quantified how sequencing depth affected genome assembly. Again, for low C_q_ samples, only 10,000 reads were required to reach fewer than 20 ambiguous bases, and with 15,000 reads (approximately 7.5 Mbp), fewer than 10 ambiguous bases remained in the genomes for all samples (**Fig. 5**).

**Figure 5.**
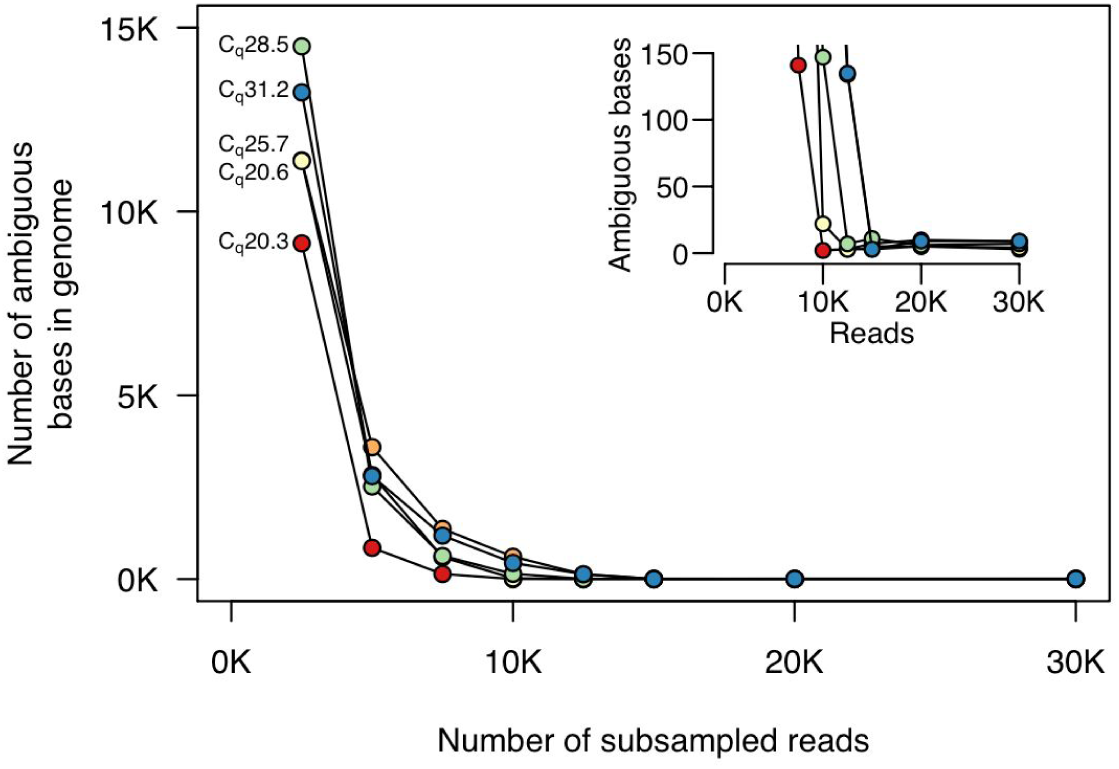
Numbers of ambiguous bases at different sequencing depths. We subsampled reads and used the filtering and assembly steps of the ARTIC Network bioinformatics pipeline. For all samples, fewer than 10 ambiguous bases remain after subsampling to 15,000 reads. For samples with lower C_q_, only 10,000 reads are required. The inset plot shows higher resolution at the lower end of the y-axis. The colours of each sample on these plots are the same as those in **Figs. 3** and **4**.

The method we use here relies on the Rapid Barcode library prep (RBK-004), which is transposon based, and usually results in only a single barcode being present on a read. This contrasts with the ligation-based library prep (LSK-109), for which most reads contain two barcodes (one at each end of the molecule). For the ligation-based library prep, stringent demultiplexing can be imposed that requires the same barcode to be present at both ends. However, when demultiplexing samples prepared using the Rapid Barcoding method, such stringent demultiplexing is not possible. Thus, there may be some crossover between samples with different barcodes due either to (1) residual transposase activity after samples are pooled; (2) low rates of ligation or chimaera formation between molecules (we observe between 0.05% and 0.1% of all reads as being longer than the longest amplicon, although many of these are human transcripts); or (3) mis-classification during the demultiplexing step performed by the basecaller and demultiplexer, Guppy. We note that the latter phenomenon is unlikely to result in high levels of barcode crossover: we find generally that fewer than 0.1% of all reads are assigned to barcodes that have not been used in any specific experiment. For this reason, we tested how read contamination affected variant calling.

We first subsampled 20,000 reads from all samples and called variants. In all cases, we found that the variant calls matched those when using all reads (between 45,000 and 115,000 for all samples (**Table 2**)). In order to analyze the effects of read contamination, we first removed sample AKL-MU-2 (C_q_ 20.6) from consideration, as the variants in this sample were identical to those in AKL-MU-5 (C_q_ 31.2). We then simulated read contamination for all pairwise combinations of samples. Each of the four samples could be contaminated by any of the other three, for a total of 12 possible combinations. The four samples we considered contained between 7 and 10 variants (**Table 2**). Sample AKL-MU-5 (7 variants) and sample AKL-MU-4 (10 variants) shared no variants with any other sample. Samples AKL-MU-1 (8 variants) and AKL-MU-3 (9 variants) shared 4 variants. Thus for only two of the 12 combinations were any variants shared. For each combination of read sets, we simulated at 27 different levels of contamination by adding between 40 and 10,000 reads from the other sample, keeping the total number of reads constant (20,000). Thus, the level of read contamination varied from 0.2% to 50%. For all combinations of read sets and contaminant levels, we called variants using the ARTIC Network bioinformatic pipeline, and tested whether these matched the variant calls when using the uncontaminated read sets.

In no cases did we find any false positive SNPs (variants that were called despite not being present in the uncontaminated original sample). Unsurprisingly, we found that as we increased contamination levels, variants were no longer reliably called. However, in all cases this only occurred at contamination levels greater than 5% (**Fig. 6**). In many cases all expected variants were called even when read contamination levels were in excess of 25%.

**Figure 6.**
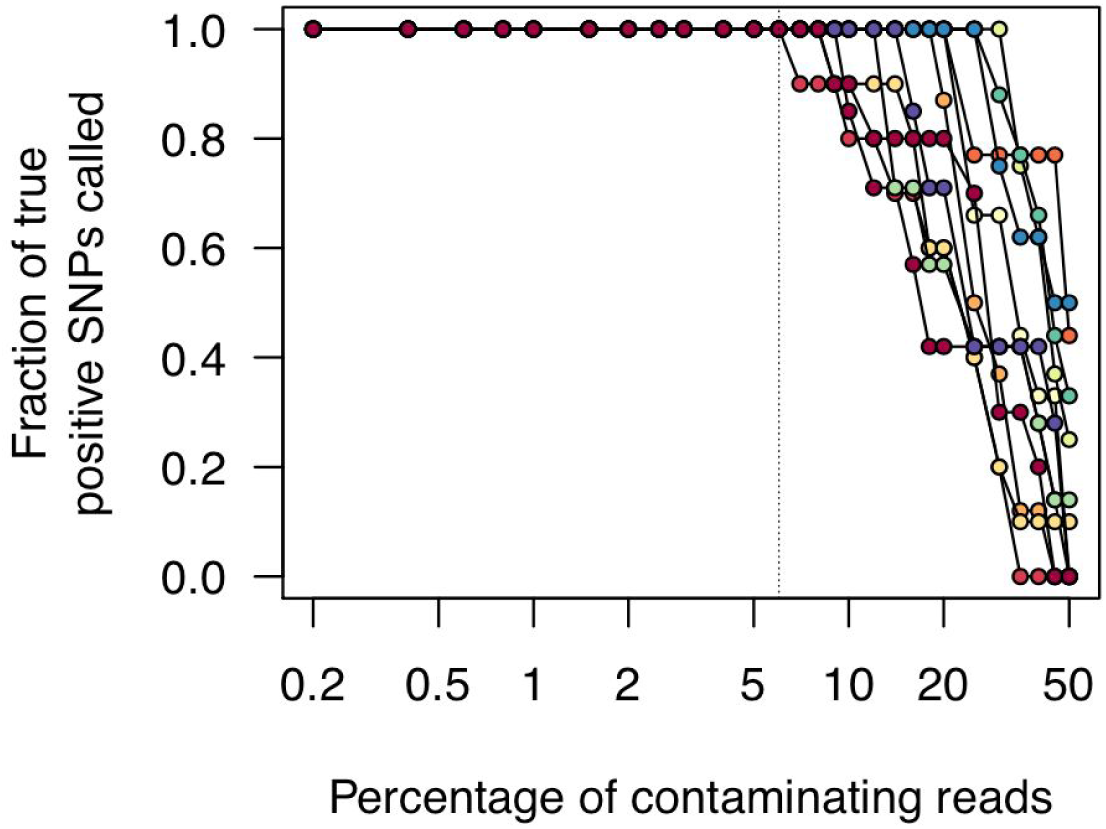
Effects of read contamination on SNP call rate. We simulated read contamination by mixing reads between all pairwise combinations of samples (see main text). We then calculated the fraction of true positive SNP calls from these contaminated read sets. Note that the x-axis is on a log scale.

## Discussion

Here we have shown that with a 1200 bp multiplexed amplicon set and the Oxford Nanopore Rapid Barcode kit we can achieve rapid, simple, and inexpensive genome sequencing of SARS-CoV-2. The sequence data results in even genome coverage, allowing high-quality viral genomes to be assembled at 50X coverage with fewer than 20,000 reads, or approximately 10 Mbp of sequencing data, and often far less. Furthermore, we have shown that even when there is read contamination, we can successfully call variants at levels of cross-sample contamination exceeding 5%. This suggests that the increased level of barcode crossover that might result when using the Rapid Barcode library prep method should rarely result in incorrect variant calls.

It will be critical to test this method on high C_q_ samples, as it is likely that achieving even genome coverage will be more difficult in these cases. Here we have only tested samples up to C_q_ 31. However, positive SARS-CoV-2 positive samples regularly have C_q_ values that are close to 37 or 38, thus having approximately 100-fold fewer genome copies per mL of sample.

This method can readily be expanded to the full set of 12 Oxford Nanopore Rapid barcodes, rather than the five we show here. There is a small amount of sequence data required for high quality genome assemblies (we estimate approximately 300 Mbp for 12 barcodes, accounting for variation in read counts among barcodes), and minimal effects of barcode crossover on variant calling. This suggests that it should be possible to sequence multiple such libraries on a single flow cell, with wash steps between sequencing runs.

## Funding

This work was funded by the New Zealand Health Research Council and Ministry of Health 2020 COVID-19 New Zealand Rapid Response Research RFP, HRC Reference: 20/1041, awarded to NF and OS.

## Acknowledgments

We thank Fahimeh Rahnama and Saed Miri-Nargesi at the Auckland District Health board for assistance in obtaining patient samples. We also thank Joep de Ligt and Una Ren for fruitful discussions.

## Data Availability

All sequence data is available at NCBI SRA XXXXXX.

## Ethics Statement

De-identified nasopharyngeal samples testing positive for SARS-CoV-2 by reverse-transcriptase quantitative (RT-q)PCR were obtained from the Auckland District Health Board Virology Laboratory. All samples were de-identified before receipt by the study investigators. This work has been provisionally approved under emergency authorisation by the New Zealand Northern B COVID-19 Human Health and Disability Ethics Committees, application number 20/NTB/85.

## Contributions

OKS and NEF conceived and designed the experiments. NEF, MV, and MBF performed the experiments. OKS performed the computational. OKS and NEF wrote the paper, with input from MV and MBF.

## Supplementary Tables and Files

**Table S1.**
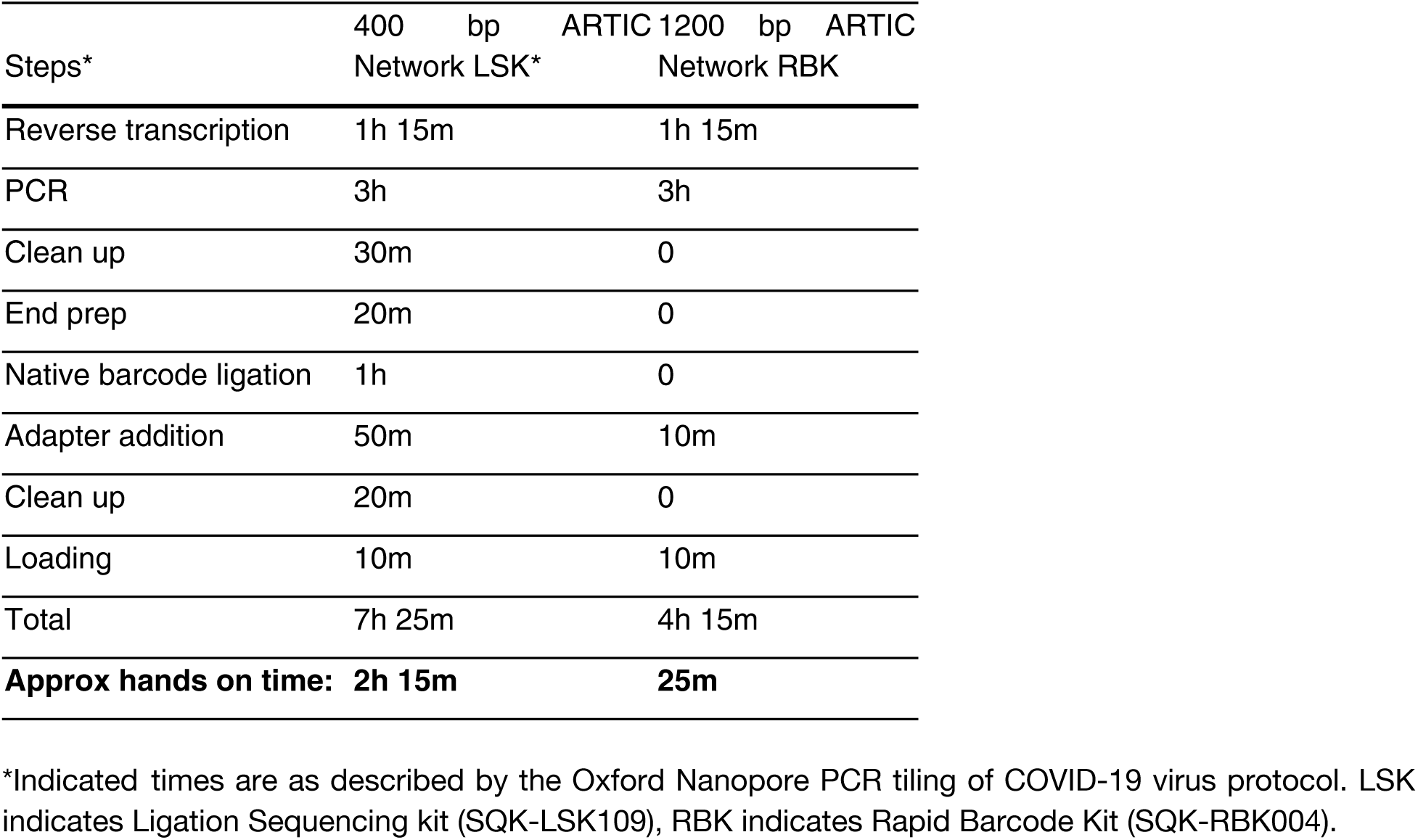
Breakdown of time to complete each step of each protocol.

## Notes

### Competing Interest Statement

The authors have declared no competing interest.

### Summary of Updates

fix sample names and title

